# Chromosomal genome assemblies of Antarctic *Notothenia coriiceps* and temperate relative *Paranotothenia angustata*

**DOI:** 10.1101/2025.10.11.681823

**Authors:** Julia M. York, Shriram Bhat, Jinmu Kim, Leyla Cárdenas, Chi-Hing Christina Cheng

**Affiliations:** Department of Evolution, Ecology, and Behavior, University of Illinois Urbana-Champaign, Urbana, IL, USA; Instituto de Ciencias Ambientales y Evolutivas, Facultad de Ciencias, Universidad Austral de Chile, Valdivia, Chile

## Abstract

Antarctic notothenioid fish genomes contain evidence of an ancient evolutionary history including their remarkable adaptive radiation beginning ∼10 MYA amid extreme selective pressures of Southern Ocean glaciation. Several Antarctic lineages subsequently escaped and recolonized temperate waters, thus their genomes chronicle the re-adaptation to warmer climates. Here, we report two chromosome-scale genomes assembled from HiFi long reads and Hi-C data for the Antarctic cryonotothenioid *Notothenia coriiceps* and its secondarily temperate close relative *Paranotothenia angustata*. The *N. coriiceps* assembly comprises 11 pseudo-chromosomes, spans 1.12 Gbp with a N90 of 81.48 Mbp and 97.8% gene complete with 24,272 annotated genes. The *P. angustata* assembly comprises 13 pseudo-chromosomes, spans 1.00 Gbp with a N90 of 62.3 Mbp and 97.9% gene complete with 23,828 annotated genes. We report two new *N. coriiceps* mitochondrial genome structures, map the antifreeze glycoprotein loci, and corroborate the unusual notothenioid ribosomal RNA gene arrangement. The high-quality assemblies of this species pair provide valuable resources for investigating both the historical evolution of polar fish and their adaptive potential to climate change.

## Background and summary

Within the perciform fish suborder Notothenioidei is a speciose group of polar fishes called cryonotothenioids that make up ∼92% of the abundance and ∼77% of the species diversity of fish in the Southern Ocean surrounding Antarctica^1^. The genome biology of this group increasingly fascinates evolutionary biologists^2^. The reasons are several: (1) Their icy Southern Ocean habitats are constantly near 0°C (range -2°C to +4°C) and frequently below the freezing point of their body fluids, necessitating the expression of antifreeze glycoproteins (AFGPs)—an early and key example of *de novo* gene birth^3^. (2) They have one of the highest rates of speciation among marine teleosts, a counterintuitive finding considering latitudinal trends in biodiversity^4^. (3) Their genomes show evidence of repeated chromosomal breaks and fusions, generating unexpectedly high karyotype diversity^5,6^ as well as mitochondrial diversity and heteroplasmy^7,8^. (4) While they are closely related, cryonotothenioids have diversified along various niche axes such as water depth and body size, to become morphologically and physiologically diverse^1,9^. (5) Some species have lost certain vital vertebrate traits such as hemoglobin, myoglobin, and the inducible heat shock response^10–12^. (6) They have undergone bursts of transposable element activity that coincide with climactic shifts^13,14^. (7) Excellent evolutionary comparisons are available from species that have secondarily re-adapted to temperate waters and variably lost Antarctic traits^15–17^.

The first notothenioid draft genome was a short-read assembly of the black rockcod *Notothenia coriiceps* published in 2014^18^. This genome was rapidly incorporated into global-scale comparison papers as the model genome of an Antarctic fish^19–21^, and frequently utilized as a reference assembly in experimental studies^22–24^ or in evolutionary genomics papers on polar adaptation^25–29^. Since 2014, genomic sequencing and annotation technologies have vastly improved, particularly with the development of highly accurate long reads that sharply reduce algorithmic collapsing of repetitive regions^30^. For genomes rich in repetitive elements including those of cryonotothenioid fishes, long reads are necessary to reveal high-level genome structure as well as accurate maps of repetitive functional regions such as the *afgp* locus^14,31^.

Here we report two chromosome level genomic assemblies for *N. coriiceps* and its closely related temperate relative *Paranotothenia angustata* (formerly *Notothenia angustata*). *N. coriiceps* is a red-blooded, demersal fish commonly found in the shelf waters of the Antarctic peninsula^32,33^. *P. angustata* (the Maori chief) diverged an estimated 10.3 million years ago and is currently found inshore along the coast of New Zealand, and reportedly also off the coast of Chile^34,35^ (divergence estimate from timetree.org). The *P. angustata* genome is the first for the species. The *N. coriiceps* assembly reported here vastly improves on the first-generation assembly; it is ∼77% larger with an assembled size of 1,124.8 Mbp consistent with the genome size estimated by flow cytometry (C-value 1.13 ± 0.21 pg)^36^, and has >3-fold greater repetitive sequence content. We annotate the complete sequence of the *afgp* region, the rDNA gene families, the mitochondrial genomes, and report on the synteny and structure of the two genomes.

## Methods

### Sampling

A male *Notothenia coriiceps* (33 cm total length; Figure 1A) was captured by baited trap January 12, 2024 at 18:20 local time off the dock of the Chilean Antarctic military Base Bernardo O’Higgins (-63.32083117163223, -57.90017358649133). The water temperature was -0.1°C. The fish was transferred to an aerated cooler and taken to the lab inside the base, where it was anesthetized with MS-222 (1 g per 15 L seawater) until opercular movement stopped (8 min 10 sec), then euthanized by cervical dislocation. Approximately 1 mL of blood was sampled from the caudal vein with a heparinized needle and syringe, immediately mixed with 50 µL heparin in phosphate buffered saline (PBS; 2000 units heparin/mL), and centrifuged at ∼10,000 x g for three minutes. The plasma was removed by pipette and replaced with an equal volume of a preservation solution (SCD: 320 mM sucrose, 40 mM sodium citrate, 5% DMSO, 1 mM EDTA, pH 7.5-8.0). The cells in SCD were immediately mixed by hand inversion several times until dispersed and then frozen and stored at -80°C until use. The fish was captured, sampled, and transported under Instituto Antártico Chileno permit 4/2024 (project FD_01-15: Centro de lnvestigación: Dinámica de Ecosistemas marinos de Altas Latitudes (IDEAL; RP2): Bio-invasions and endemism: a physiological and molecular approach in the Southern Ocean).

**Figure 1:**
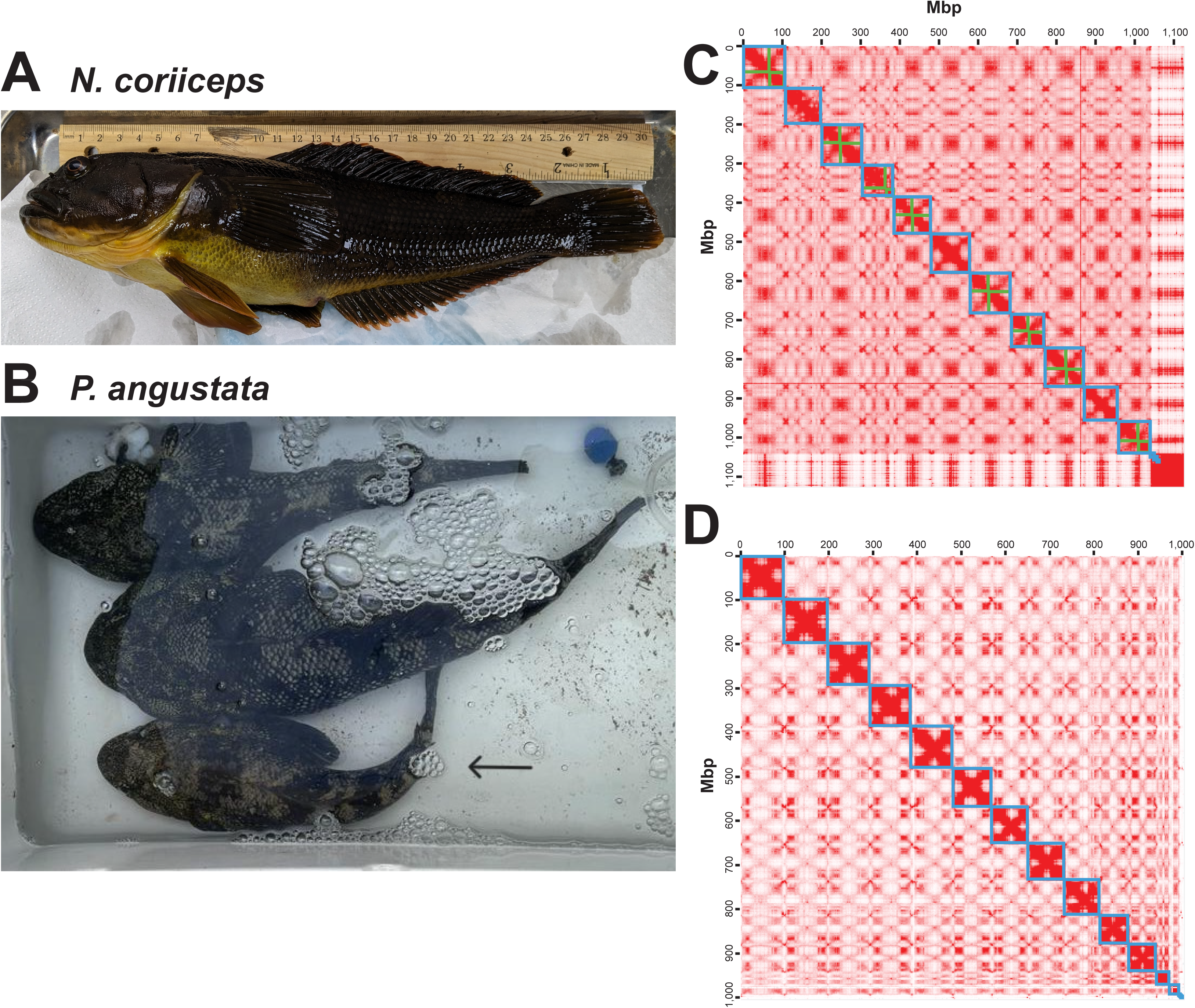
(A) *Notothenia coriiceps* specimen sampled for this study. (B) *Paranotothenia angustata* specimens sampled (genome individual indicated by arrow). Hi-C contact maps for (C) *N. coriiceps* and (D) *P. angustata* with scaffolds show in blue boxes. Green boxes on *N. coriiceps* contact map indicates boundaries of chromosomal arm mis-joins, which were manually corrected to be consistent with linkage maps as described supplemental table 4 (detail on 1C lower right in supplemental figure 3).

An immature female *Paranotothenia angustata* (33 cm total length; Figure 1B) was captured by baited trap on August 7, 2023 at 13:09 local time near the Shag Point Matakaea picnic area on the South Island of New Zealand (-45.4684, 170.828095). The water temperature was 9.54°C. The fish was kept in an aerated cooler at the field site, anesthetized with MS-222 (250 mg per L seawater) until opercular stop (9 min 13 sec), and euthanized by cervical dislocation. A blood sample was taken, centrifuged, and the erythrocytes were preserved in SCD as above, flash frozen in liquid N_2_ and then stored at -80°C. Both fish species were captured, held, and euthanized in accordance with University of Illinois Urbana-Champaign IACUC approved protocol 23104.

### DNA isolation and sequencing

About 50 – 100 uL of SCD preserved frozen blood cells were scraped from sample tubes into 1 mL ice cold PBS and pelleted at 0°C by centrifuging for 5 minutes at 6000 x g. Supernatant was removed, and cells were resuspended in 200 µL ice cold PBS. High molecular weight (HMW) DNA was extracted using the Nanobind PanDNA kit (PacBio 103-260-000) following PacBio protocol (102-574-000) for nucleated red blood cells. DNA size and integrity were assessed on a 1% agarose gel, and concentrations were determined using the Qubit dsDNA Broad Range assay with a Qubit 3 fluorometer (Invitrogen). The HMW DNA was submitted for PacBio HiFi library construction at the Roy G. Carver Biotechnology Center, University of Illinois Urbana-Champaign. The DNA integrity and molecular weight were re-confirmed with Femto Pulse (Agilent). HMW DNA was sheared with a Megaruptor 3 (Diagenode) to a target average fragment length of 15 kbp, which was then made into a HiFi sequencing library using the SMRTBell Prep kit 3.0. The libraries from the two species were two-plexed and sequenced on one PacBio Revio SMRT cell 8M for 30 hours of data capture, which generated 38.5 Gbp sequence data for *N. coriiceps* and 25.6 Gbp for *P. angustata*. To augment the coverage for *P. angustata*, a separate fraction of the HiFi library was sequenced on another half Revio SMRT cell, which generated an additional 21 Gbp of sequence data, totaling 46.6 Gbp for the species. Hi-C scaffolding libraries were prepared from liver and spleen by Phase Genomics (Seattle, WA) using their Proximo platform, which entailed a restriction enzyme (DpnII, DdeI, Hinfl, and MseI) digest of the cross-linked chromatin. The Hi-C libraries were sequenced with 150 bp paired-reads on an Illumina NovaSeq X Plus 10B lane, generating 186.4 million read pairs for *N. coriiceps* and 118.5 million read pairs for *P. angustata*. To increase read coverage for *P. angustata*, another aliquot of its Hi-C library was sequenced, generating an additional 65.3 million read pairs and totaling 183.8 million for the species.

### RNA-Seq

All existing RNA-Seq data available for *N. coriiceps* on the National Center for Biotechnology Information (NCBI) website was downloaded from the sequence read archive (SRA) to use for genome annotation (see supplemental table 1). The sequences were derived from tissues obtained in various experimental studies: heart, brain, gill, and liver (SRP470441); skin, head kidney, and duodenum (SRP367384); liver, brain, blood, and skin (SRP031878); as well as head kidney expression data (SRP069032) used to annotate the first *N. coriiceps* genome^18^. No existing RNA-Seq data was publicly available for *P. angustata* at the time of this study. We sampled heart, whole brain, liver, spleen, pyloric ceca, stomach, pancreatic tissue, eye, caudal kidney, head kidney, intestine, red and white muscle, ovary, and pectoral fin from two euthanized individuals dissected on ice (see supplemental table 2 for specifics). Tissues were stored in 90% ethanol at -20°C, replacing the ethanol after 24 hours. To isolate RNA, tissues were bead homogenized with 0.5 mm zirconium oxide beads in TRIzol (Invitrogen) using a Bullet Blender (Next Advance). Total RNA was extracted following Invitrogen protocol for TRIzol, then digested with DNaseI and column purified with the Monarch RNA cleanup kit (New England Biolabs). RNA was checked for quality on a 1% denaturing agarose gel and for concentration with the Qubit RNA Broad Range assay kit. Equal amounts (0.75 µg) of RNA from each tissue replicate were pooled, and a library was prepared using the WatchMaker mRNA prep kit (WatchMaker Genomics) with poly A+ selection. The library was sequenced on an Illumina NovaSeq X Plus 10B lane generating a total of 481 million 150 bp paired end reads.

### Assembly

We used *`hifiasm`* in Hi-C integration mode (version 0.16.0)^37^ to assemble the HiFi and Hi-C reads into the primary assembly which was used in all downstream analyses. Contigs were checked for contamination and adaptor content with *`FCS-GX`* and *`FCS-adaptor`* (version 0.5.5)^38^. Initial Hi-C scaffolding of the *N. coriiceps* genome produced 12 chromosomal-size scaffolds accounting for 94% of the assembled length, as opposed to the n=11 haploid chromosome number known for the species^6^. The smallest scaffold 12 (∼31 Mbp), contained only 5 gene models, low repeat content, ∼3.4 Mbp region rich in rDNA sequences, and shared no clear unique homology to any region in the other 11 large scaffolds (supplemental figure 6). A large rDNA region in the unplaced scaffold 12 suggested that *`hifiasm`* might have assembled allelic rDNA genes as distinct copies and failed to place the resulting scaffold into a chromosome. We therefore ran *`purge_dups`* (version 1.2.5)^39^ on the primary Hi-C integrated assembly followed by Hi-C scaffolding, which eliminated this unplaced scaffold 12. Both assemblies with and without *`purge_dups`* are available in Illinois Data Bank repository (IDB-7296574). The *P. angustata* assembly did not have *`purge_dups`* applied. For Hi-C scaffolding, mate pair reads were aligned to the primary assembly with *`bwa-mem2`* (version 2.2.1)^40^, PCR duplicates were marked and removed with *`SAMBLASTER`* (version 0.1.26)^41^, and the contigs were scaffolded with *`YaHS`* (version 1.2)^42^. The Hi-C contact map was generated with *`Juicer`* (version 1.19.02)^43^ and visualized with *`Juicebox`* (version 2.15; Figure 1C and D)^44^. The mitochondrial genomes were assembled from the raw HiFi reads using *`MitoHiFi`* (version 3.2.2)^45^, using the *N. coriiceps* RefSeq mitochondrial genome (NC_015653.1) as a reference.

### Manual editing

We utilized published linkage maps for a female and male *N. coriiceps* constructed using RAD-seq to refine the chromosome assignment of our assemblies^46^. We aligned the RAD-tag sequences from the linkage maps to the scaffolded assembly of both species and visualized the synteny using *`macrosyntR`* (version 0.2.19; supplemental figure 1)^47^. *P. angustata* has 13 chromosomes (2n=26)^5^ hypothesized to be a result of 11 pairwise fusions of the ancestral n=24 karyotype, an unfused chromosome 12, and a single pericentromeric inversion of chromosome 13. Each *N. coriiceps* linkage group cleanly mapped to a single scaffold in the *P. angustata* assembly, except for linkage groups 2 and 4, each of which mapped to two scaffolds as predicted by the karyotype fusion hypothesis (supplemental figure 1)^46^. We assigned chromosome numbers to *P. angustata* scaffolds to match the numbering of the homologous linkage groups (supplemental table 3 and Figure 2).

**Figure 2:**
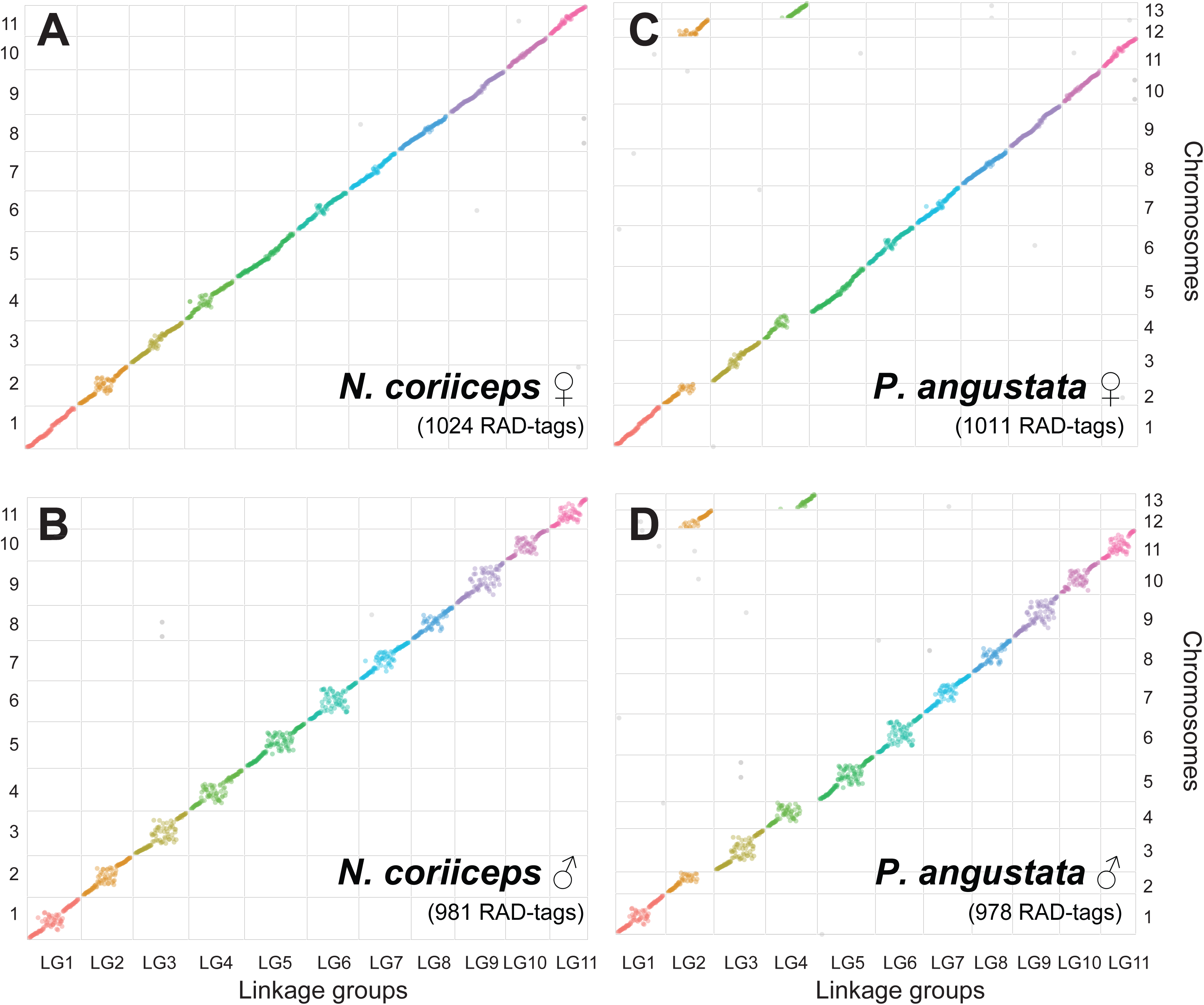
Synteny with *N. coriiceps* linkage maps after manual editing. Locations of RAD-tag sequences in the assemblies (y-axes) vs the linkage maps (x-axes) from Amores et al.^46^ (A) *N. coriiceps* female, (B) *N. coriiceps* male, (C) *P. angustata* female, and (D) *P. angustata* male. Note that each linkage group corresponds to a single chromosome in the *N. coriiceps* assembly but LG2 and LG4 are split in two chromosomes each for *P. angustata*, consistent with the hypothesis that these chromosomes subsequently fused to create the *N. coriiceps* 2n=22 karyotype.

The 11 haploid chromosomes of *N. coriiceps*^6^ was deduced to result from two additional fusions after divergence from *P. angustata*, namely chromosome 2 fused with chromosome 12 and chromosome 4 fused with chromosome 13^6,46^. The linkage map RAD-tags did not map as cleanly to our initial Hi-C scaffolded *N. coriiceps* assembly, indicating some high-level assembly inconsistencies. Guided by the published *N. coriiceps* linkage maps and evidence obtained from telomere sequence localization detailed below, we manually edited the Hi-C scaffolds to arrive at the final ordered chromosome set (see supplemental table 4). Specifically, linkage maps and karyotype suggested that all 11 chromosomes are approximately 100 Mbp in length^46^, however, our initial Hi-C scaffolded assembly recovered one unusually long scaffold of nearly 200 Mbp (scaffold 1; supplemental figure 1). We reasoned that this scaffold resulted from two chromosomes mis-joined by the assembler, as we found linkage group 8 mapped to the first half of the scaffold and linkage group 9 mapped to the second half (supplemental figure 1). To locate the potential mis-join, we used *`Telomere Identification toolKit`* (*`tidk`*; version 0.2.64)^48^ to detect telomere sequences that typically populate chromosome termini. We compared the positions of the detected peaks of telomere repeats to the positions of the RAD-tags sequences in our assembly. In the unusually long scaffold 1, a peak of telomere repeat sequences coincided with the junction between linkage groups 8 and 9 (supplemental figure 2). This result led us to manually split scaffold 1 at the telomere peak into two separate scaffolds. Further, by referencing the RAD-tag order and orientation we identified that the two resulting scaffolds from splitting scaffold 1, as well as scaffolds 2, 3, 5, and 8, were apparently assembled with the chromosome arms inverted—such that the telomeres were joined at the center and the centromere regions from the homologous linkage groups were at the ends of these scaffolds (supplemental figure 1). Indeed, using *`tidk`*, we found telomere sequence peaks coinciding with each putative mis-join (supplemental figure 2). We therefore manually reoriented the arms of these scaffolds. In addition, scaffolds 9 and 12 mapped to each half of linkage group 4, and scaffolds 10 and 11 mapped to each half of linkage groups 3, so these scaffolds were manually joined (supplemental figure 1). All scaffolds were then assigned chromosome numbers based on linkage group homology (Figure 2). Full details on manual editing and chromosome numbering are given in supplementary table 4. Recombination rate was estimated by dividing the distance in Mbp between the most distal linkage map RAD-tags for each chromosomal scaffold by the frequency of recombination between those loci in centiMorgans, averaged across all 11 chromosomes.

### Structural analysis

Centromere locations were predicted using *`CentroMiner`* from the *`quarTeT`* package (version 1.2.5)^49^. Synteny was compared using alignments generated with *`nucmer`* from the *`mummer`* package (version 4.0.1)^50^, filtering out alignments that were less than 50% unique using the delta-filter utility (-u 50), and generating plots using *`circos`* (version 0.69.8)^51^. Synteny plots with the platyfish *Xiphorphorus maculatus* were generated similarly (assembly GCF_002775205.1_X_maculatus-5.0-male). Assemblies were assessed for contiguity with *`QUAST`* (v.5.2.0)^52^ and completeness with *`BUSCO`* (v.5.8.1; odb12)^53^.

### Repeat content

To explore the repeat content we ran comprehensive transposable element annotator *`Earl Grey`* (version 6.3.2)^54^. We provided *`RepeatMasker`* (version 4.1.8; repeatmasker.org) within *`Earl Grey`* with a *de novo* repeat library for each genome generated with *`RepeatModeler2`* (version 2.0.6)^55^ combined with the root (0) and Eupercaria (10) partitions of the dfam library (version 3.9; dfam.org).

### Annotation

RNASeq reads were trimmed and quality checked using *`fastp`* (version 0.24.1)^56^, *`fastqc`* (version 0.12.1)^57^, and *`multiqc`* (version 1.28)^58^, then aligned to assemblies using *`HISAT2`* (version 2.2.1)^59^, which achieved a 94.2% overall alignment rate for *N. coriiceps* and 96.8% overall alignment rate for *P. angustata*. Aligned RNASeq reads and repeat masked assemblies were provided to *`BRAKER3`* for structural annotation (version 3.0.8)^60^. Protein functional annotation was conducted using *`InterProScan`* (version 5.70-102.0)^61^ and *`AGAT`* (version 0.7.0; github.com/NBISweden/AGAT). Assemblies were scanned for ncRNA using *`infernal`* and the Rfam 15 database (version 1.1.5)^62,63^.

### AFGP and rDNA annotation

The highly repetitive coding sequences of *afgp* genes elude detection as coding genes by automated annotation tools such as *`BRAKER3`*, requiring manual annotation. The approximate positions of the *afgp* copies were first mapped by blast searches of the assemblies using published *afgp* gene sequences (HQ447059.1; HQ447060.1)^64^. The *afgp* genomic region was then extracted from the assembly and the structure of each member gene was manually annotated using the visualization and mapping tools in *`SnapGene`* (version 8). We also analyzed the nuclear rDNA gene families (5S, 5.8S, 18S, and 28S) by manually mapping and annotating the tandem arrays of member genes using the same approach. The rDNA queries were retrieved from the RefSeq genome of the non-Antarctic notothenioid *Eleginops maclovinus*, the closest sister taxon of the Antarctic cryonotothenioids (GCF_036324505.1)^65^. The *`blastn`* matches for each species were used as queries for a second species-specific *`blast`* search to ensure all copies were found. We scored the full-length matches as valid rDNA copies.

## Data record

### Sequencing

*N. coriiceps* PacBio HiFi sequencing generated 38.6 billion base pairs in 2.96 million reads, with an average length of 13,021 bp (range 97-59,775). The genome was scaffolded using 186.4 million Hi-C read pairs and annotated with the 534.4 billion base pairs of publicly available RNASeq reads on NCBI (Figure 1C and supplemental figure 3). The *P. angustata* genome was assembled from 46.6 billion base pairs across 3.33 million reads averaging 13,980 bp in length (range 88-56,261), Hi-C scaffolded using 183.8 million read pairs, and annotated using 481 million bp in newly generated RNASeq reads (Figure 1D). All raw data and assemblies are available on NCBI (Bioproject PRJNA1310647).

### Assemblies

The final *N. coriiceps* assembly is 1.12 Gbp in length, 92.6% of which is found in 11 chromosomes (karyotype 2n=22; Table 1; Figure 1C), and 97.8% gene complete according to BUSCO (Table 1). The genome was sequenced to an average depth of 45x, and 99.4% of the assembly had coverage >10x. No contamination was detected. The *N. coriiceps* genome is 56.7% repetitive elements: 10.9% DNA transposons, 13.6% retrotransposons, 8.5% simple and other repeats, and 23.7% unclassified repeats (supplemental figure 4). The published linkage map for this species found 11 linkage groups each approximately 103 cM in length^46^. Using the probabilities of recombination from the linkage maps we estimate the male recombination rate of *N. coriiceps* to be 1% for every 975,461 ± 26,111 bp or 1.03 ± 0.03 cM/Mbp.

**Table 1:**
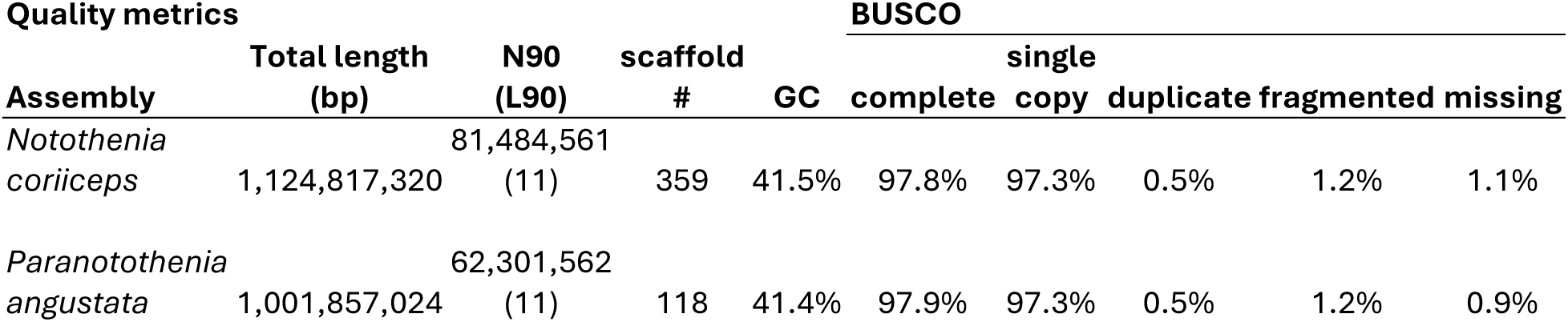
Assembly statistics.

The *P. angustata* assembly is 1.00 Gbp in length, 99.1% of which is found in 13 chromosomes (karyotype 2n=26; Table 1; Figure 1D). The average sequencing coverage depth for the entire assembly was 54x, for the 13 chromosomal scaffolds it was 35x, and 99.56% of the assembly had coverage >10x. No contamination was detected. The assembly is 52.1% repeats: 10.4% DNA transposons, 14.3% RNA transposons, 4.8% simple or other repeats, and 22.6% unclassified repeats (supplemental figure 5). Overall, the macrosynteny between the two species is highly conserved (Figure 3).

**Figure 3:**
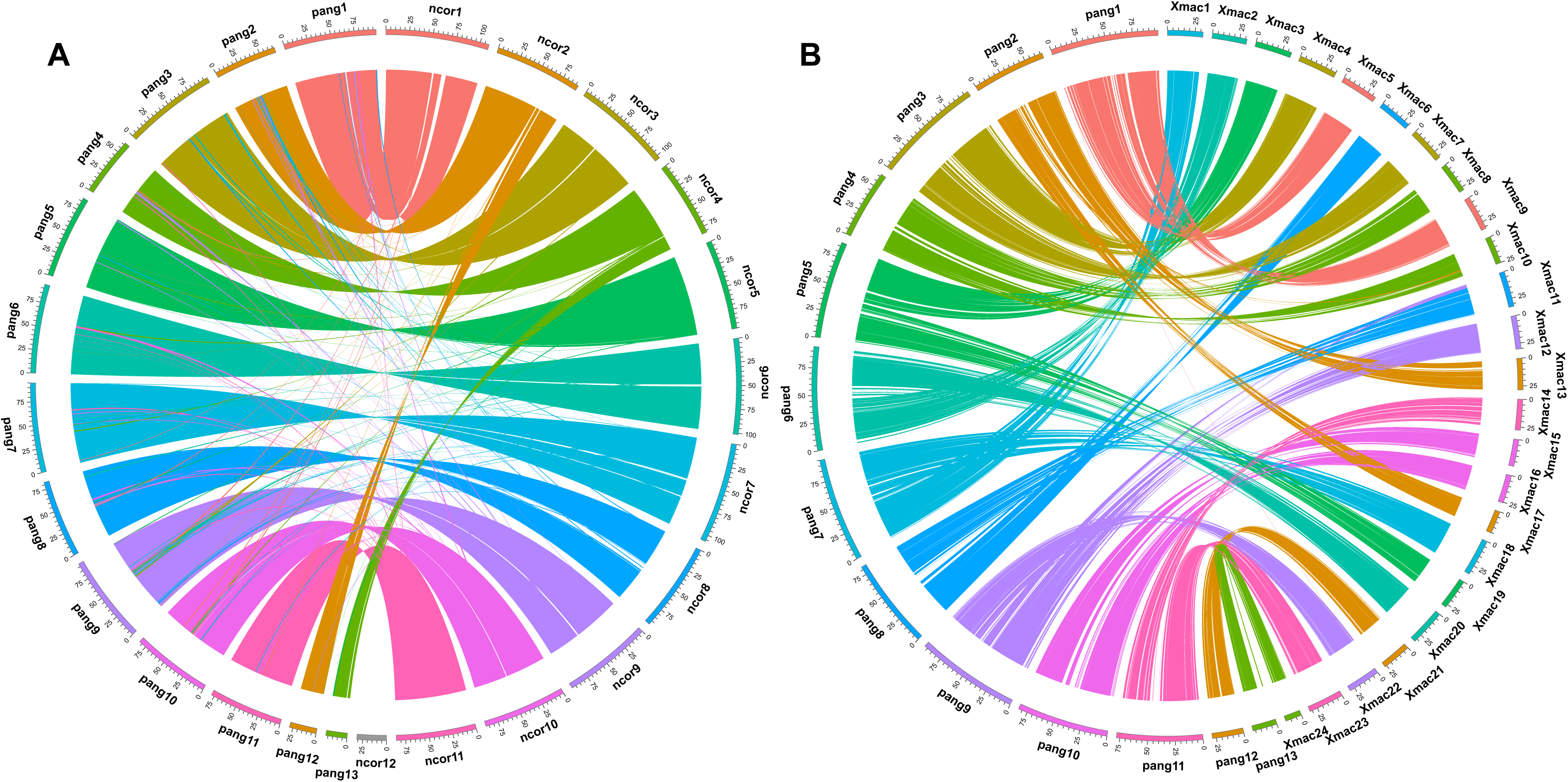
Macrosynteny between (A) the assembled chromosomes *N. coriiceps* and *P. angustata* and (B) *P. angustata* and *Xiphophorus maculatus* indicating alignments between assemblies that are at least 50% unique matches.

### Annotation

A total of 24,272 genes were structurally annotated for *N. coriiceps*, and 96.5% of these were functionally annotated. The coding sequences represented 5.3% of the total genome length (Figure 4A). For *P. angustata*, 23,828 genes were structurally annotated, 97.0% of which were functionally annotated, and the coding sequences made up 5.7% of the genome (Figure 4B).

**Figure 4:**
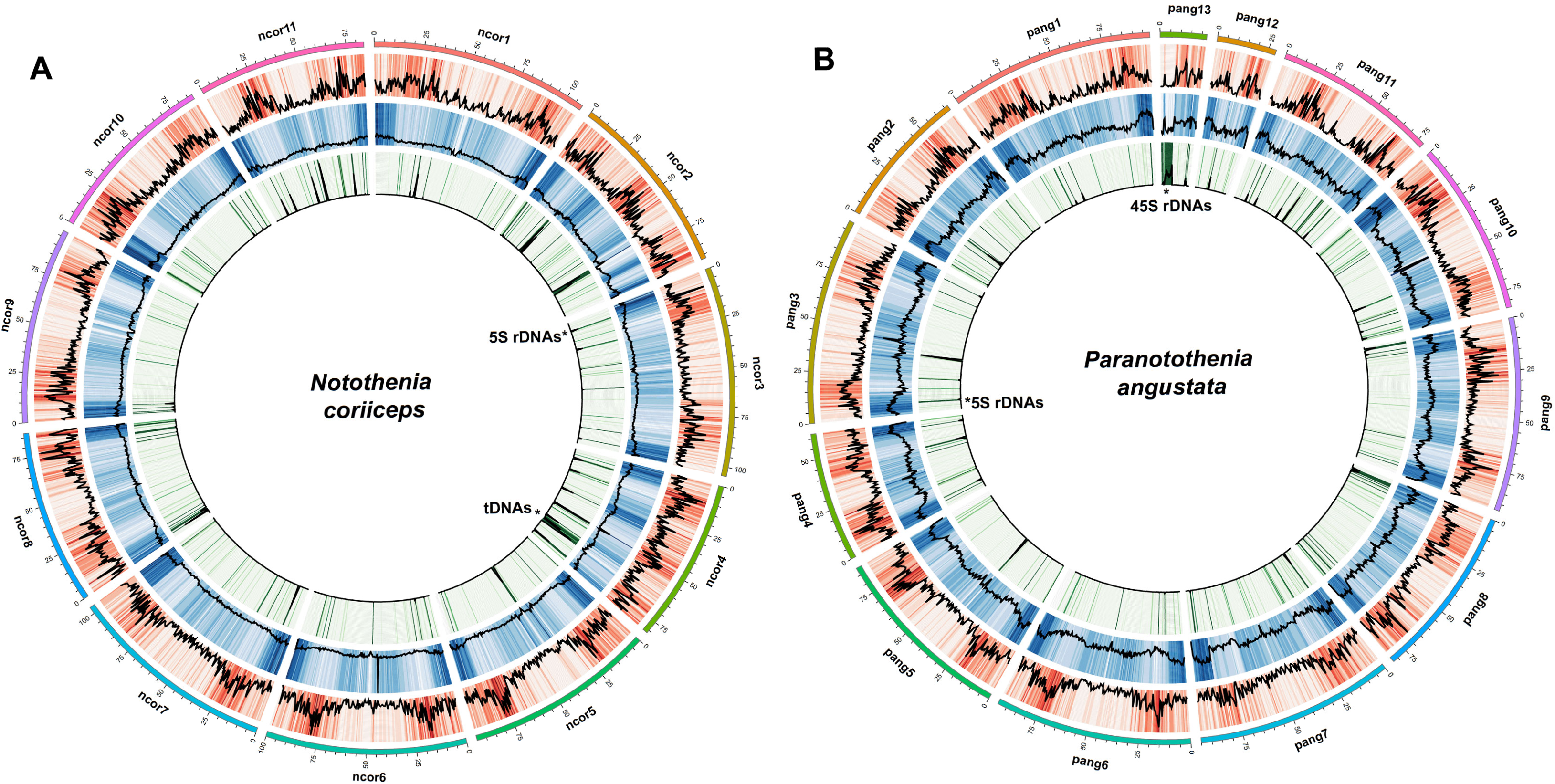
Chromosomal structures of (A) *N. coriiceps* and (B) *P. angustata* assemblies. Chromosomes are indicated by outer colored bands (ncor1, pang1… etc.), ticks indicate sequence lengths in Mbp. Outer red ring shows density of protein coding gene models, middle blue ring density of repetitive elements, and inner green ring density of DNA coding for ncRNA. 5S and 45S rDNA regions indicated were confirmed by manual annotation.

### Mitochondrial genomes

Two contigs from each species contained structurally distinct mitochondrial genomes. For *N. coriiceps*, neither contig matched any published gene order (Figure 5A)^8,66^. However, they contain rearrangements known to occur in cryonotothenioid fishes, namely the translocation of *nad6* and adjacent tRNA for Glu and Pro into the control region (CR) and as well as the varying degree of duplication of these elements and the CR. One contig contains two duplicates of CR1 (atg000001l; NCBI accession PX233742). The other contains three copies of CR1, three copies of tRNA-Pro, as well as two duplicates of *nad6,* one of which was pseudogenized (*nad6*ψ) by a 1 bp reading frame shift (ptg000001c; accession PX233743). The occurrence of two distinct mitochondrial genome structures indicates mitochondrial heteroplasmy in *N. coriiceps*; this inference is supported by adequate sequence coverage, ranging from 5-22x for atg000001l and 6-10x for ptg000001c.

**Figure 5:**
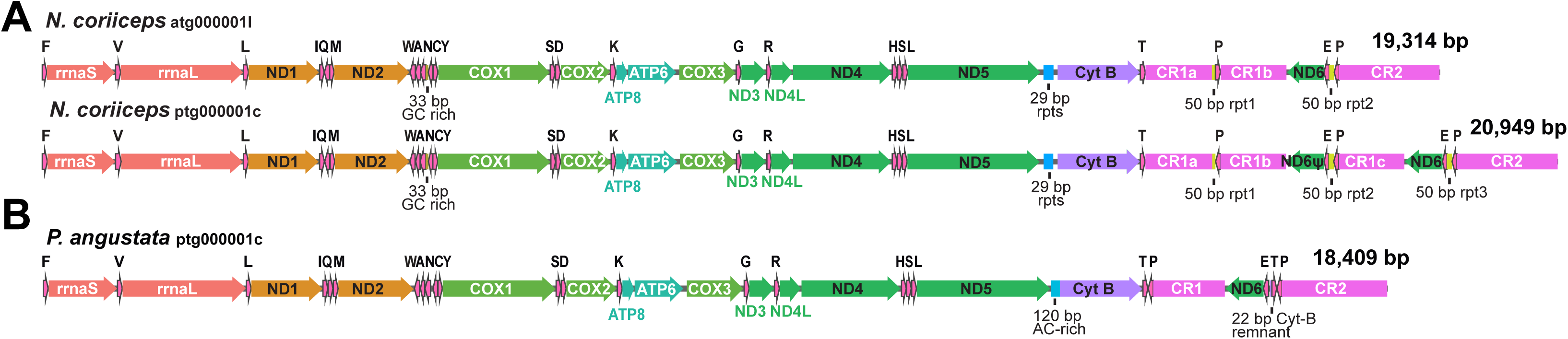
Mitochondrial assembly gene orders. One contig is shown for *P. angustata* (A) and two were found with similar read coverage for *N. coriiceps* (B). Arrows indicate gene order and direction. Control region duplications are labeled with letters (CR1a, CR1b… *etc*).

For *P. angustata*, one contig had high coverage (20-40x) and matched the Noto1GO order described by Papetti et al.^8^ with the putative tandem duplication of the *nad6* through CR1 region followed by partial random loss (ptg000001c; NCBI accession PX233744; Figure 5B). The other contig (ptg000002l) lacked CR1, *nad6*, and tRNAs for glutamic acid and proline, but had low read coverage, particularly in this putative loss region (1-2x coverage) and therefore was not considered reliable.

### Antifreeze glycoproteins

The *afgp* genomic region spans 575 kbp in *N. coriiceps*, of which 339 kbp comprises the large *afgp* locus, encompassing 17 copies of *afgp* and one chimera of *afgp* and *tlp* (trypsinogen-like protease). All copies except *afgp*10ψ have intact gene structures and are presumed functional (supplemental table 6). In *afgp*10ψ, there is a single in-frame premature stop codon leading to a shorter AFGP polyprotein precursor, however this gene could still produce 17 AFGP molecules after translational processing. Collectively, the *N. coriiceps afgp* gene family can generate a total of 612 AFGP molecules spanning all size isoforms in a single round of transcription and translation (Figure 6 and supplemental table 6).

**Figure 6:**
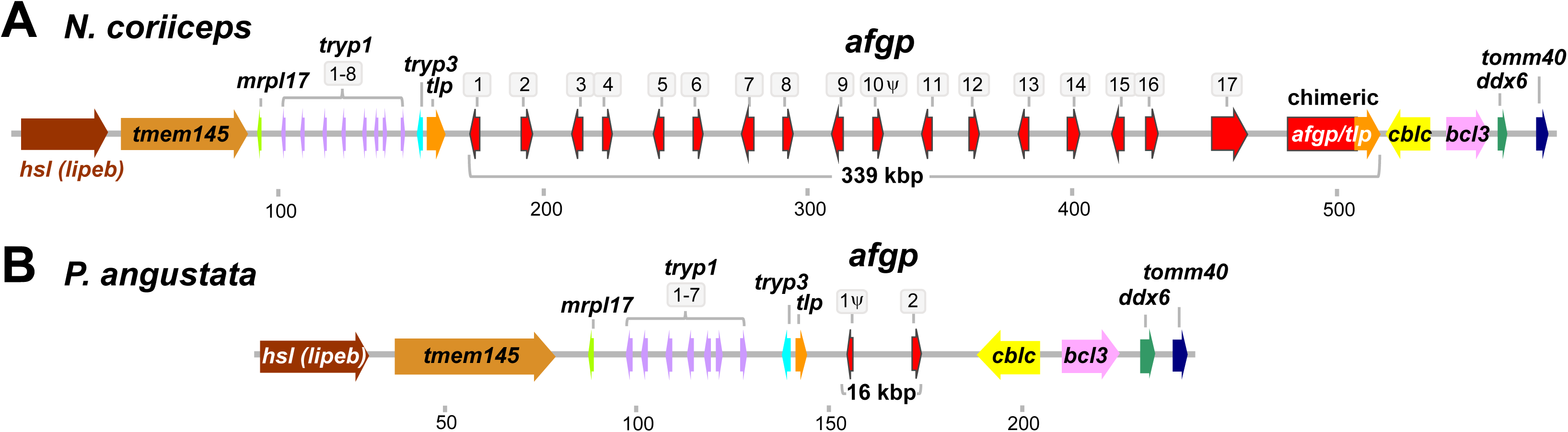
The AFGP locus of (A) *N. coriiceps* and (B) *P. angustata*. Gene location and direction are indicated by arrows, outlined arrows indicate *afgp* copies. Scale is in kbp.

The secondarily temperate New Zealand relative *P. angustata* has a residual *afgp* locus of just 16 kbp consisting of only two copies of *afgp.* Copy 1 lacks exon 1 that codes for the signal peptide and is thus nonfunctional. Copy 2 appears structurally intact and it encodes 18 AFGP molecules of the smallest isoform, but these contain amino acid substitutions that likely render the proteins inactive (Figure 6; supplemental table 7).

### Ribosomal RNA genes

The major 45S rDNA loci for both species consisted of tandemly repeated units of 5S-28S-5.8S-18S, with the 5S preceding 28S in the opposite direction (Figure 7). For *P. angustata*, the major locus assembled on chromosome 13 and was ∼1.3 Mbp in length with 32 tandem 45S units. A separate ∼91 kbp locus containing only 5S rDNA was found on chromosome 3 (Figure 4). For *N. coriiceps*, the homologous 5S locus was accurately assembled in the same region of chromosome 3, spanning a 161.8 kbp region (Figures 4 and 7). However, the 45S locus did not assemble in the expected location on chromosome 4 (the homolog of *P. angustata* chromosome 13). Instead, a dense cluster of 60 tandem 5S-28S-5.8S-18S units spanning 1.93 Mbp were assembled at one end of the unplaced scaffold 12 of the *N. coriiceps* genome assembled without *`purge_dups`* (Figure 7 and supplemental figure 6). Applying *`purge_dups`* before re-scaffolding the primary assembly was intended to remove potential rDNA duplicates and improve 45S rDNA locus assembly. However, it did not lead to successful recovery and placement of the locus in chromosome 4, rather, clusters of the 45S rDNA units remained in multiple small unplaced scaffolds underscoring that accurate assembly of long tandem arrays of repeating genes remains a bioinformatics challenge^30,67^.

**Figure 7:**
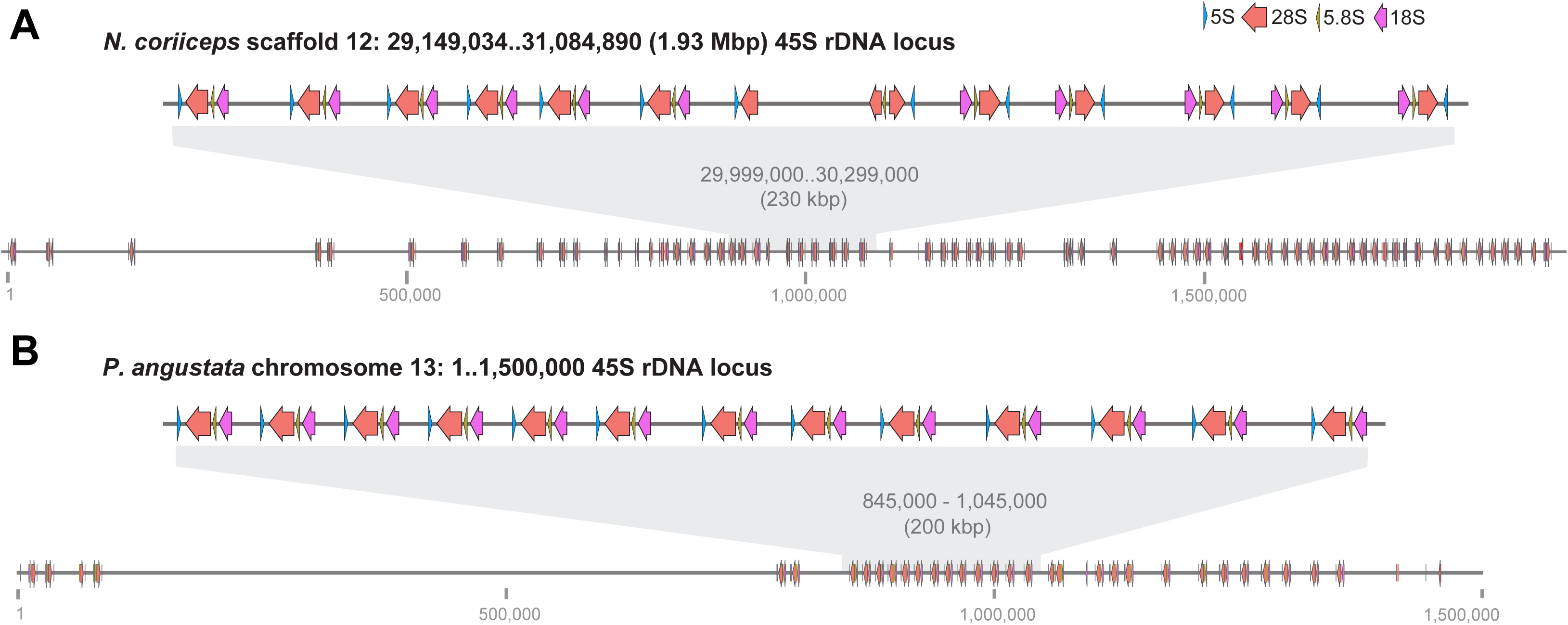
The major nucleolar organization region (NOR) rDNA loci in (A) scaffold 12 of the *N. coriiceps* assembly before additional haplotig purging with *`purge_dups`* (see supplemental figure 6); the most rDNA-dense region is shown (∼1.93 Mbp out of ∼3.4 Mbp), and (B) a ∼1.3 Mbp region of chromosome 13 of the *P. angustata* assembly. A 230 kbp and 200 kbp section, respectively, is expanded to show gene order of the tandem 45S units that are each proceeded by 5S rRNA gene copies in the opposite orientation. Gene location and direction are indicated by arrows, gene identity by color, scale is in bp.

## Technical validation

Our genome assemblies corroborate the chromosomal fusion hypothesis of Amores and colleagues^46^ which posits that the n=24 ancestral teleost karyotype, best exemplified among model teleosts by *Xiphophorus maculatus*, underwent massive pairwise fusion of 22 chromosomes and a single pericentromeric inversion to generate the n=13 karyotype of *P. angustata* (Figure 3). Subsequently, ancestral chromosomes 12 and 13 fused with chromosomes 2 and 4, respectively, to generate the n=11 karyotype of *N. coriiceps*. Our two assemblies resolve these chromosomes, including telomeres and centromeres (supplemental figure 2) with 93% and 99% of the assembly lengths found within the chromosomal sequences.

The quality of the *N. coriiceps* assembly reported here improves significantly on the 2014 assembly, which was smaller (636.6 Mbp), with an L90 of >13,000, 18.2% repeats, and 77% complete BUSCOs^18^. Our long-read assembly recovered >3-fold more repetitive content than the 2014 short-read assembly likely due to the algorithmic collapsing of repeats during the assembly process. The total length of our newly generated *N. coriiceps* assembly matches the genome size expected from previous analyses (estimated 1.11 Gbp; Table 2)^36^.

**Table 2:** Chromosome lengths.

| chromosome | *N. coriiceps*<br>length (bp) | *P. angustata*<br>length (bp) |
| --- | --- | --- |
| 1 | 108,698,235 | 98,251,549 |
| 2 | 92,061,790 | 66,820,301 |
| 3 | 104,489,007 | 99,291,504 |
| 4 | 81,484,561 | 62,301,562 |
| 5 | 94,772,574 | 88,675,529 |
| 6 | 100,116,035 | 94,355,478 |
| 7 | 103,467,516 | 95,089,416 |
| 8 | 87,149,981 | 82,687,813 |
| 9 | 97,840,480 | 93,162,105 |
| 10 | 87,593,405 | 83,381,870 |
| 11 | 83,907,317 | 78,225,433 |
| 12 |  | 28,627,820 |
| 13 |  | 22,315,419 |

We estimate the recombination rate for the male *N. coriiceps* to be 1.03 ± 0.03 cM/Mbp, comparable to the 1.13 cM/Mbp estimated in humans, but very low for a teleost fish which more typically have recombination rates around 2 cM/Mbp^68,69^.

Two mitochondrial genomes were assembled for *N. coriiceps* with reasonably high coverage, but neither matched the published notothenioid gene orders. Cryonotothenioids are known to have high levels of diversity in mitochondrial genome structure and heteroplasmy^7,8^. This adds an additional gene order to the more than 10 already described for the group, an extremely high level of mitochondrial structural diversity given the relatively young age of the clade^8,14^. The *P. angustata* gene order matched that of the most basal cryonotothenioid mitochondrial structure (Noto1GO) from Papetti and colleagues^8^. Another mitochondrial genome contig for *P. angustata* assembled but lacked *nad6* and tRNA-Glu, a mutation which was previously described^70^ and then refuted^66^. However, this contig had low coverage especially in the region lacking *nad6* and tRNA-Glu, and so we considered it unreliable.

The novel antifreeze glycoprotein (*afgp*) is a hallmark of Antarctic notothenioid evolutionary adaptation to the freezing Southern Ocean. Notothenioid *afgp* genes encode large AFGP polyprotein precursors, thus the coding sequences are long runs of 9-bp (3-codon) repeats encoding the constituent conserved tripeptides (Thr-Pro/Ala-Ala), resembling simple sequence repeats. These highly repetitive sequences almost invariably elude recognition as protein-coding in automated annotation, necessitating manual detection. In the *P. angustata* assembly, only a remnant remains of the *afgp* locus, with one nonfunctionalized copy of *afgp* and the other with multiple amino acid substitutions. In contrast, the *afgp* locus of Antarctic *N. coriiceps* is >20-fold larger and the translated polyproteins could produce 612 AFGP molecules spanning all size isoforms in a single round of transcription and translation. These findings are consistent with studies on the respective antifreeze protein expression levels in these two species^71,72^.

Despite major progress in genome sequencing technologies, tandem arrays of rDNA remain one of the most difficult regions to assemble with fidelity^30,67^. The eukaryote 45S rDNA locus typically comprises tandem repeats of the 28S-5.8S-18S gene module, forming the nucleolar organization region (NOR). The presence of 5S rDNA genes upstream of the 28S in both the species studied here corroborates previous findings suggesting this ribosomal RNA gene structure is a synapomorphy of cryonotothenioids and unique thus far among fishes^73^. In curating bait rDNA sequences from the basal sister notothenioid *E. maclovinus*, we discovered that the 5S-28S-5.8S-18S structure also exists in this basal non-Antarctic relative, thus we conclude this trait is not a synapomorphy of the Antarctic notothenioid clade alone (supplemental figure 7).

The non-NOR 5S loci are found in homologous locations on chromosome 3 of each species (Figure 4 and supplemental figure 8). However, the major 45S locus is currently unresolved in *N. coriiceps*, despite cytogenetics data demonstrating a single large 45S NOR in this species^74^. In *P. angustata*, even when a 45S locus partially assembled on chromosome 13, smaller clusters are still found in various unplaced scaffolds. We expect the 45S locus in *N. coriiceps* to localize in chromosome 4—the homolog of the 45S locus-bearing chromosome 13 of *P. angustata*. Future work improving repeat assembly methodology may help to resolve this region and uncover the evolutionary dynamics of the ribosomal RNA gene families^75^.

## Usage notes

The *P. angustata* genome is an excellent assembly for understanding climate adaptation, as this lineage is of Antarctic ancestry but later re-colonized temperate waters. For example, multiple Antarctic marine species have lost the otherwise universal heat shock response (HSR) characterized by inducible heat shock protein expression^76,77^. *Bovichtus variegatus*, a basal notothenioid that never adapted to Antarctic waters retains the HSR, while *P. angustata* continues to lack a functional HSR despite ∼10 million years away from the Antarctic^78^. Further study utilizing this genome could deduce whether the regulatory or coding sequences necessary for the HSR have been degraded or lost in *P. angustata* and why the HSR trait has not been recovered. Similarly, Antarctic notothenioids typically show high ubiquitination of their proteins, thought to aid in coping with cold denaturation, and *P. angustata* appears to retain this trait and thus is phenotypically similar to the Antarctic group^79^. The characterizations of the rDNA gene families are foundational for future inquiries into the evolution of rRNA genes across both Antarctic and non-Antarctic notothenioids, which appear to share a non-canonical 45S rDNA structure seen otherwise only in crocodilians and testudines^73^. Additionally, in at least one cryonotothenioid species rDNA loci for 5S and 45S have expanded onto two chromosomes each^75^. We report these assemblies and annotations as a resource to delve into such questions and continue the study of intriguing questions around Antarctic fish genome evolution.

## Data availability

The raw sequencing reads (.fastq), primary genome assemblies (.fasta), and newly generated RNASeq data (.fastq) are available on NCBI (Bioproject PRJNA1310647). Mitochondrial sequences can be found on NCBI accessions PX233742-PX233744. Antifreeze glycoprotein and rDNA mapping details can be found in the supplemental. Annotations (gff3, cds, protein), the unpurged *N. coriiceps* assembly, and other supporting files can be found on Illinois Data Bank (repository DOI: 10.13012/B2IDB-7296574_V1).

## Code availability

No custom code was generated; all bioinformatics tools were used with default parameters except where noted.

## Acknowledgements

The authors would like to thank Captain Edwin Vidal and the logistics team at Base O’Higgins, Emilio Alarcón, Sebastián Brauchi, Ellen York, Macgregor Willis, and Alvaro Gonzalo Hernandez and his sequencing team at the Roy G. Carver Biotechnology Center at the University of Illinois Urbana-Champaign.

## Author contributions

CHCC and JMY conceived of the project; CHCC, LC, and JMY collected and sampled the specimens; CHCC oversaw the sequencing efforts; JMY, CHCC, SB, and JK conducted the analysis; JMY, CHCC, and SB prepared the figures; JMY and CHCC prepared the manuscript; and SB, JK, and LC contributed to manuscript editing.

