## supplemental for "Chromosomal genome assemblies of Antarctic *Notothenia coriiceps* and temperate relative *Paranotothenia angustata*"

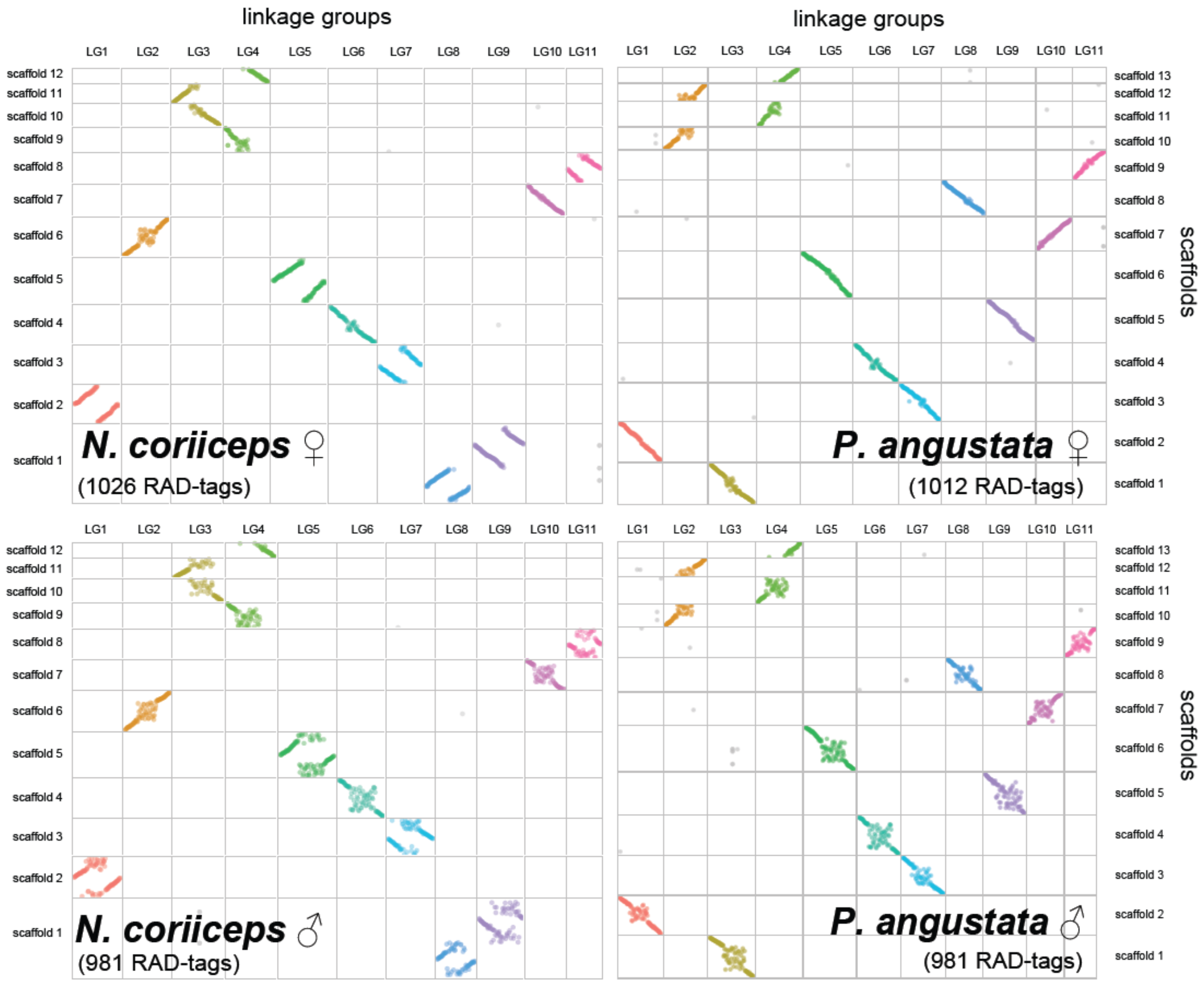


Supplemental figure 1: synteny of the linkage group RAD-tags from Amores et al.[46] with the scaffolded assemblies before manual editing. Note that the linkage group RAD-tags map linearly to scaffolds in the *P. angustata* assembly but several *N. coriiceps* scaffolds have a distinct juncture at the center where highly linked RAD-tags are found at opposing ends of the scaffolds (chromosome arms inverted and joined at the telomeres). Further, scaffold 1 of *N. coriiceps* maps distinctly to two linkage groups and linkage groups 3 and 4 are found on two scaffolds each. Manual editing details can be found in supplemental table 4.

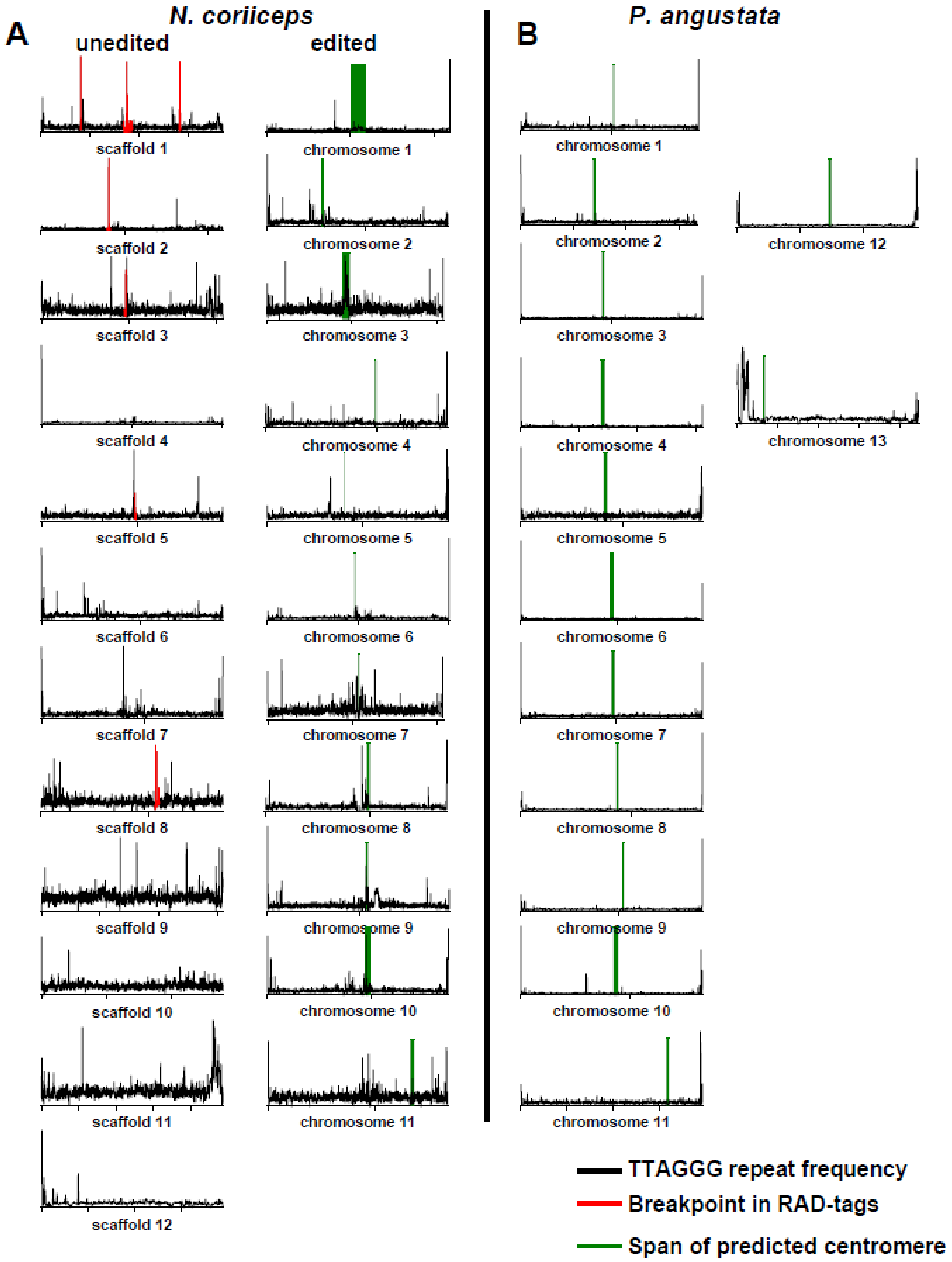


Supplemental figure 2: telomere repeat frequencies and predicted centromere locations across scaffolds and chromosomes for (A) *N. coriiceps* unedited (left) and edited (right) assemblies; (B) *P. angustata* edited assembly. Black indicates frequency of telomere repeats (TTAGGG) in either direction, green indicates predicted centromere span, and red indicates the breakpoints between linkage group RAD-tag orders and presumed mis-joins of chromosome arms that were manually corrected.

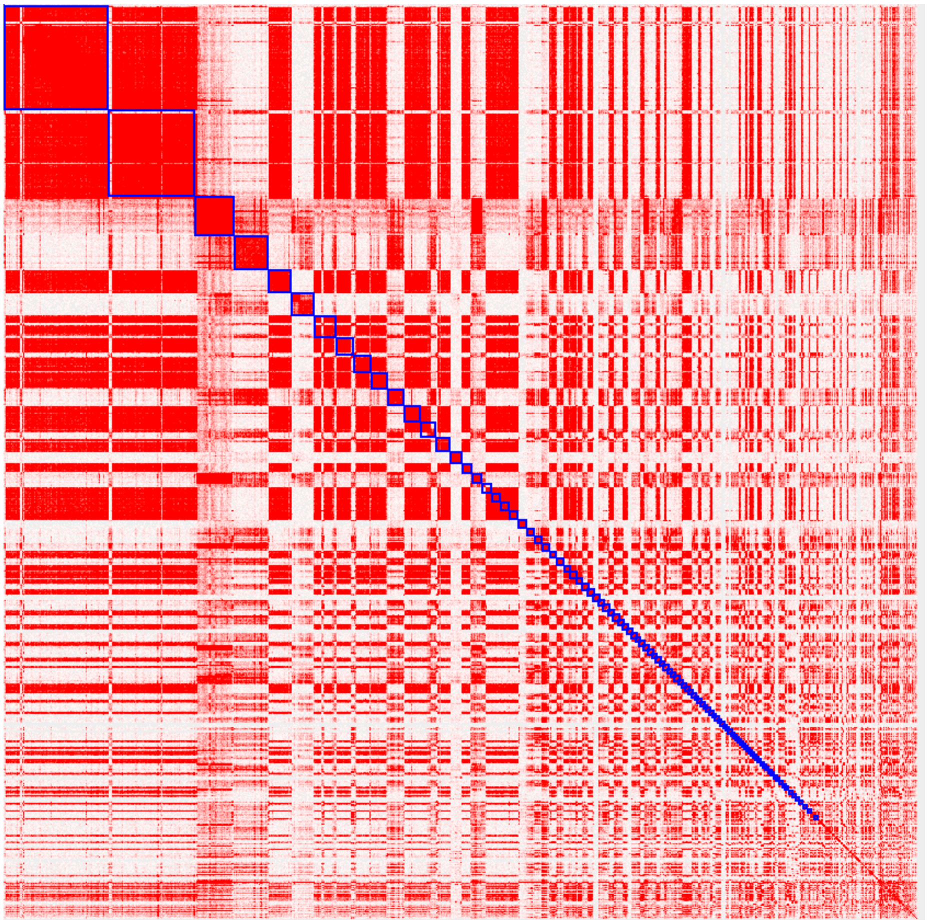

Supplemental figure 3: Hi-C contact map of the unplaced scaffolds for *N. coriiceps*.

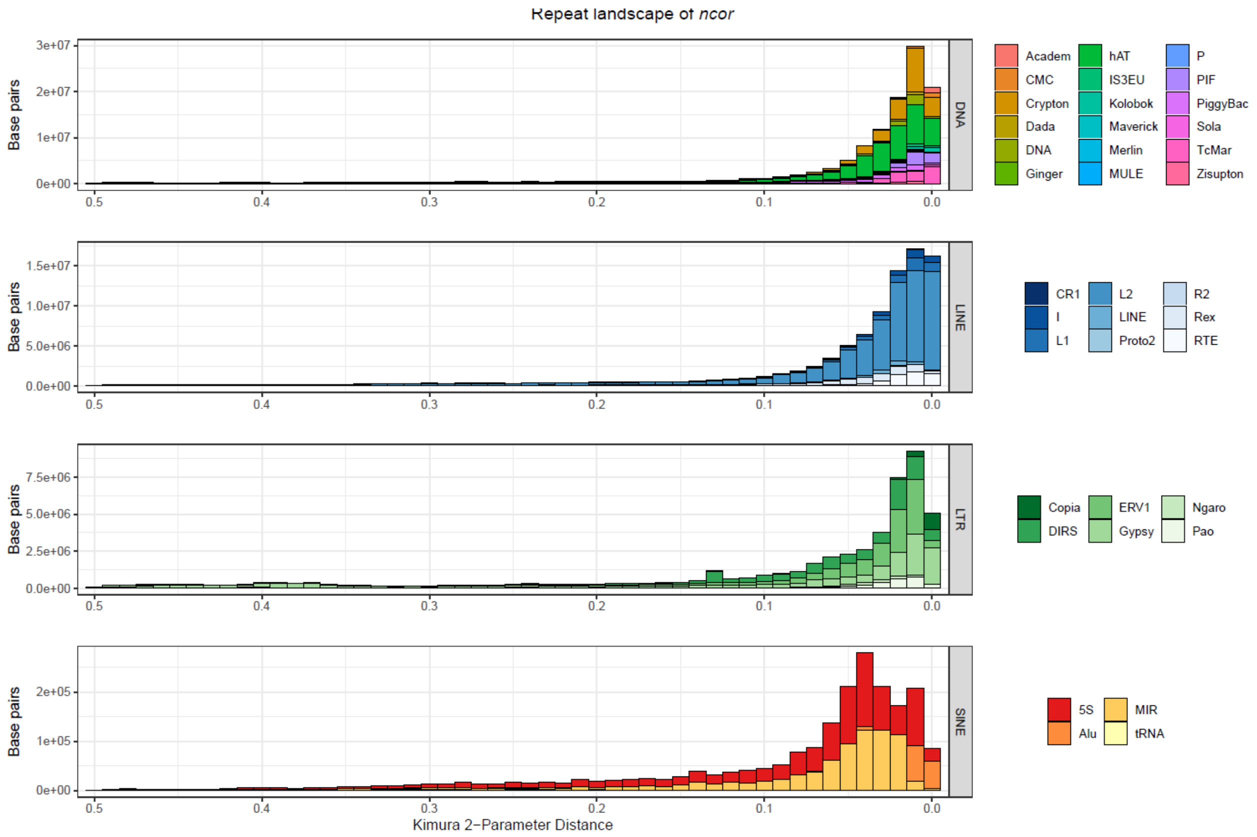


Supplemental figure 4: Kimura 2-parameter distance of repeat element superfamilies for *N. coriiceps*.

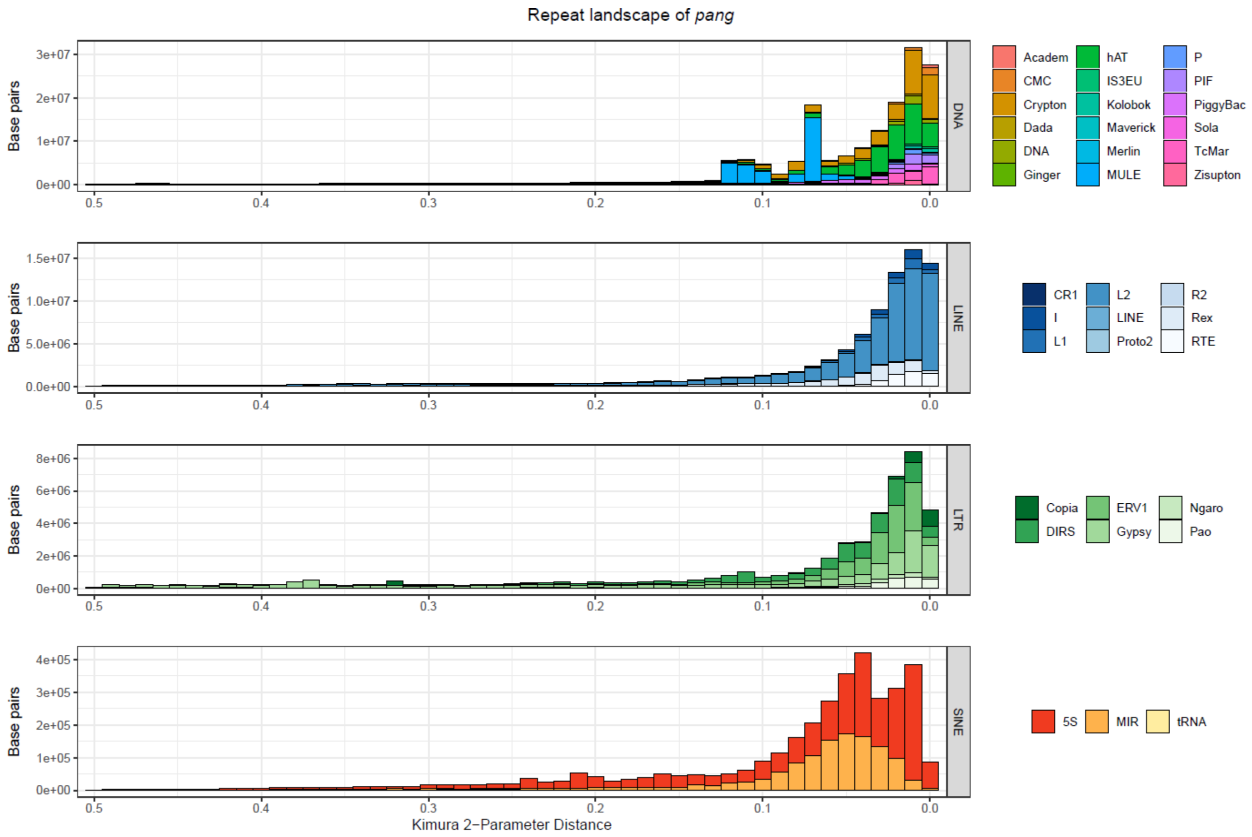


Supplemental figure 5: Kimura 2-parameter distance of repeat element superfamilies for *P. angustata*.

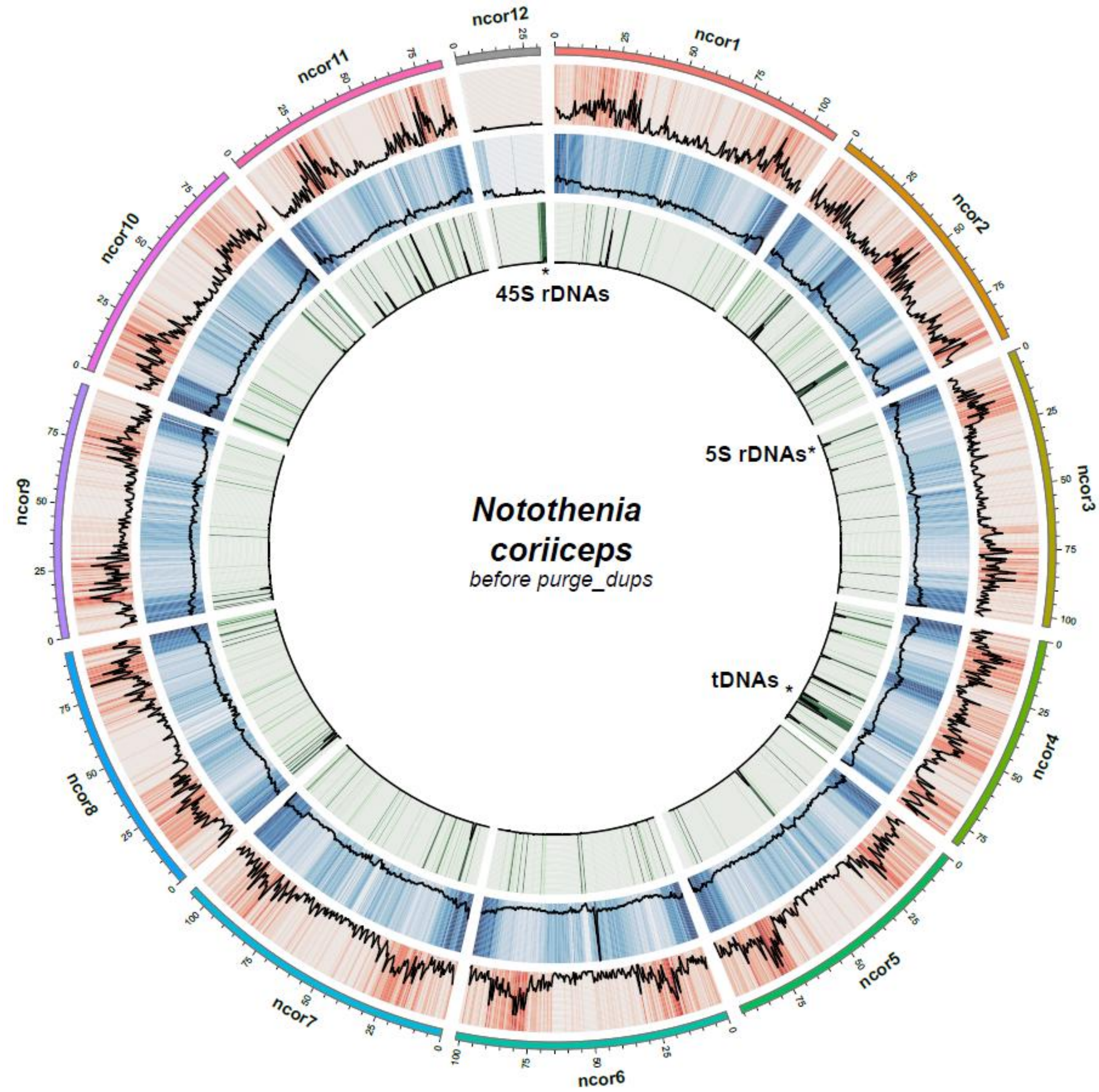


Supplemental figure 6: Gene, repeat, and DNA coding for ncRNA density (outer, middle, inner circles, respectively) for *N. coriiceps* assembly before `purge_dups` was used to purge haplotigs. Note the large density of 45S rDNA on unplaced scaffold 12.

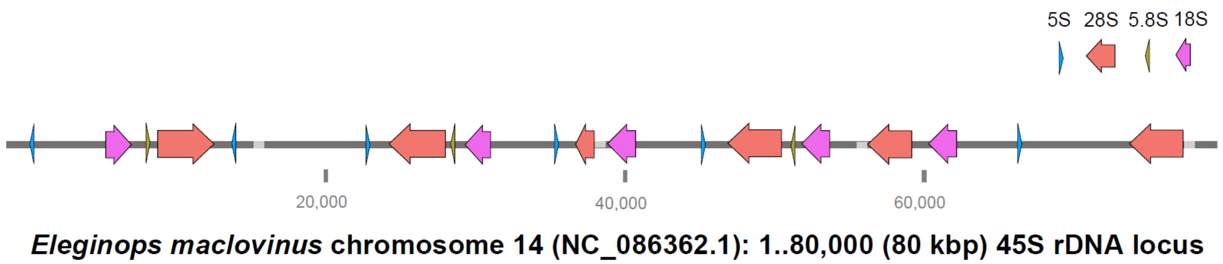


Supplemental figure 7: The 45S ribosomal RNA gene locus in chromosome 14 (accession NC_086362.1) of the genome assembly of *Eleginops maclovinus*, the closest non-Antarctic sister taxon of the Antarctic cryonotothenioids. The assembled locus spans ~80 kbp, with low rDNA gene content, and contains gaps (light grey boxes on the dark grey bar), indicating that this assembled region is likely incomplete. The rDNA content includes three 45S units with full-length 28S-5.8S-18S rRNA genes, and each unit is preceded by a copy of 5S in the opposite orientation, as was also observed in *N. coriiceps* (Antarctic) and *P. angustata* (non-Antarctic) in this study. We therefore conclude the 5S-28S-5.8S-18S structure of the 45S locus evolved before the divergence of the Antarctic clade of notothenioids. Arrows indicate the order and orientation of rRNA gene copies.

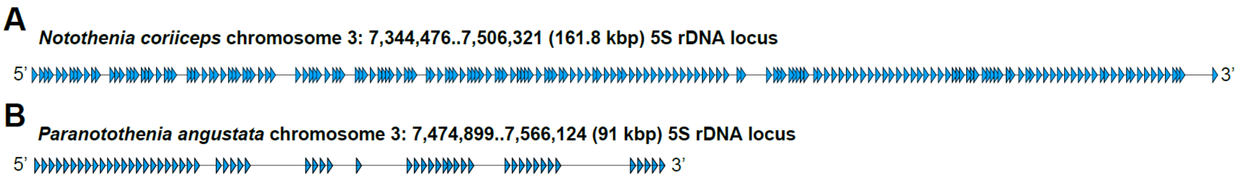


Supplemental figure 8: The 5S ribosomal RNA gene locus in chromosome 3 of (A) *N. coriiceps* and (B) *P. angustata*. The 161.8 kbp locus in *N. coriiceps* contains 182 full-length (119 bp) tandem copies predominantly spaced at 856 bp or 463 bp apart. The 91 kbp locus in *P. angustata* contains 58 full-length copies predominantly spaced at 866 bp or 878 bp apart.

Note: supplemental tables (All_Supplemental_Tables.xlsx and in .csv format) can be found on the data repository on Illinois Data Bank: https://doi.org/10.13012/B2IDB-7296574_V1

Supplemental table 1: RNASeq data used for the *BRAKER3* structural annotation of *N. coriiceps*.

Supplemental table 2: collection notes (top) and RNA extraction details (bottom) for *P. angustata*, used for *BRAKER3* structural annotation.

Supplemental table 3: manual edits and resulting chromosome numbering of the *P. angustata* assembly.

Supplemental table 4: manual editing and resulting chromosome numbering of the *N. coriiceps* assembly.

Supplemental table 5: chromosome lengths, telomere repeat counts and predicted centromere locations for each assembly.

Supplemental table 6: translated sequences of the AFGP molecules/isoforms encoded by the 17 copies of *afgp* and the single copy of chimeric *afgp/tlp* in *N. coriiceps*. "rpts" refers to the (Ala/Pro-Ala-Thr) tripeptide repeats comprising the AFGP molecule. Each encoded isoform is linked to the next by a conserved 3-residue of Leu-Ile/Asn-Phe that is post-translationally removed. A total of 612 AFGP molecules/isoforms may be synthesized in one round of transcription and translation of the 17 genes.

Supplemental table 7: translated sequences of the AFGP molecules/isoforms in the two *afgp* copies of *P. angustata*. Brown amino acids are substitutions. The green tripeptide "APT" is non-canonical.
